# Pan-cancer spatial characterization of key immune biomarkers in the tumor microenvironment

**DOI:** 10.1101/2025.03.05.641733

**Authors:** James R. Lindsay, Jennifer Altreuter, Joao V. Alessi, Jason L. Weirather, Anita Giobbie-Hurder, Ian Dryg, Katharina Hoebel, Bijaya Sharma, Kristen Felt, F. Stephen Hodi, Neal Lindeman, Lynette M. Sholl, Ethan Cerami, Jonathan A. Nowak, Mark M. Awad, Scott J. Rodig, William Lotter

## Abstract

Deciphering the composition and spatial organization of the tumor immune microenvironment (TiME) is key to uncovering the mechanisms driving cancer progression and treatment response. Spatial biology techniques like multiplex immunofluorescence (mIF) offer detailed insights into the TiME but are often limited to retrospective research studies of individual cancer types. Conversely, bulk omics techniques have been studied in pan-cancer settings but fail to capture single-cell spatial information. Here, we provide a pan-cancer spatial characterization of key biomarkers (CD8, FOXP3, PD-1, PD-L1) of the TiME using data from a mIF assay performed prospectively in a clinical setting on 2,019 tumors across 14 major cancer types. By integrating interpretable compositional and spatial metrics, we identified patterns of TiME variation that are conserved across cancer types and stages. We assess associations between these TiME spatial factors and tumor, genomic, and clinical features, where the results both extend prior findings and uncover new links. Altogether, our findings offer pan-cancer insights of the TiME to further the fields of spatial biology and cancer immunology.

## Introduction

Tumors develop in complex microenvironments containing a variety of immune and stromal cells that can both enhance and suppress cancer progression. Understanding the cellular composition and spatial organization of these tumor immune microenvironments (TiMEs) is important for developing more effective therapeutic strategies^1,2^. To this end, spatial biology techniques, such as multiplex immunofluorescence (mIF), provide detailed views of the TiME by measuring biomarkers in tissue samples while preserving spatial context^1–8^. Studies using mIF imaging have demonstrated associations between TiME features and patient outcomes^3–5,9–11^, such as the relationship between intratumoral CD8+ and FOXP3+ cell densities and prognosis in lung^5,10^ and colorectal cancer^11^. However, most mIF studies to date have focused on individual cancer types in pre-selected populations. Furthermore, important opportunities remain to explore how mIF-derived spatial features correlate with the genomic phenotypes of tumors.

Here, we characterize spatial aspects of the TiME based on a standardized mIF assay^12^ that was performed prospectively in a clinical setting on tumor tissue specimens from 2,019 tumors across over 14 cancer types (**Fig. 1**). The assay interrogates four key immune biomarkers (CD8, FOXP3, PD-1, PD-L1) as well as a tumor cell marker and DAPI for nuclei detection. By combining metrics of cellular composition and spatial organization, we first aimed to identify the major patterns of TiME variation across the cohort. We then sought to quantify how these patterns relate to tumor, genomic, and clinical features.

**Figure 1:**
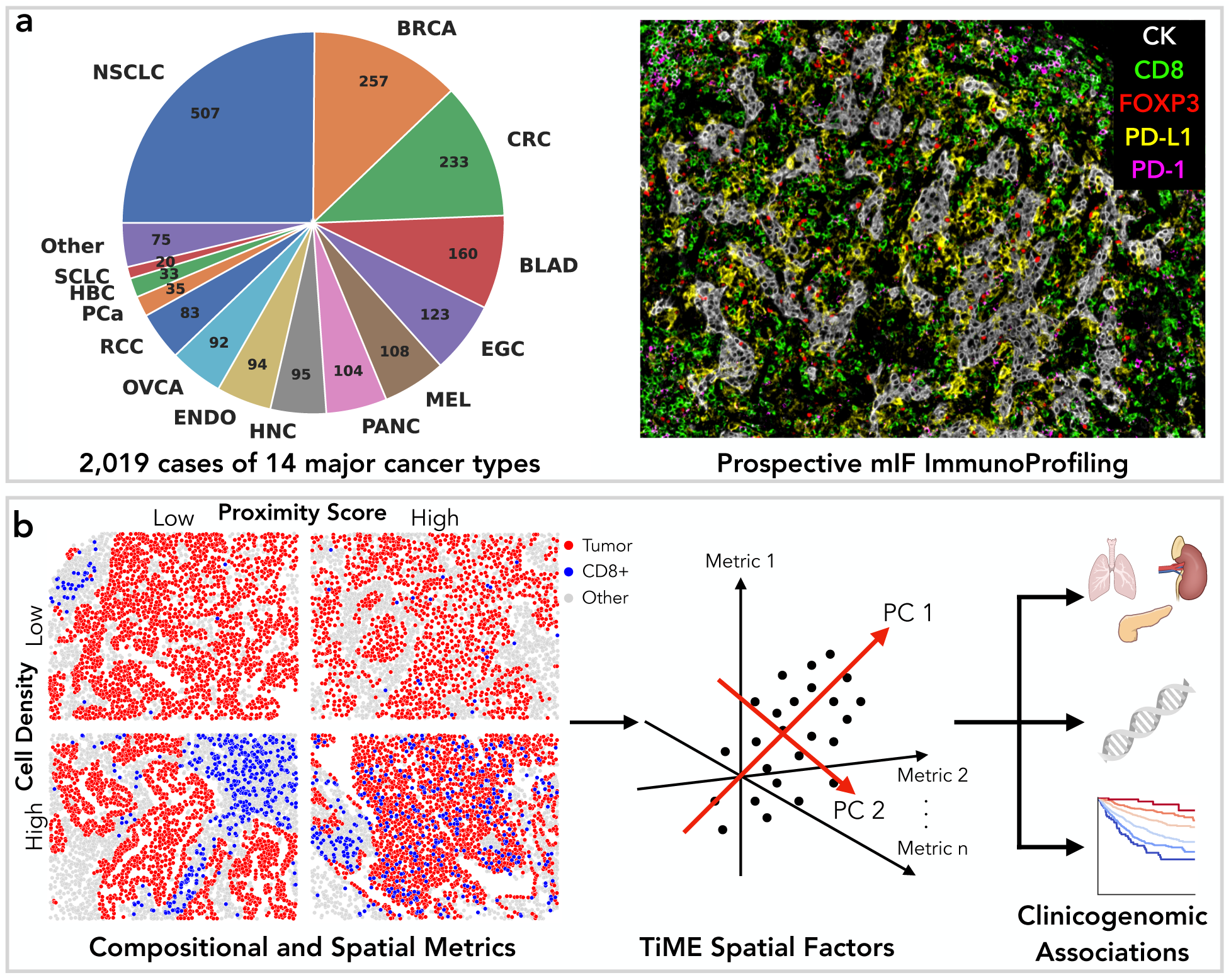
Pan-cancer spatial characterization of the TiME. a) A mIF assay (ImmunoProfile) was performed prospectively across 2,109 cases representing 14 major cancer types. The distribution of cancer types and an example ROI are displayed. b) Summary of computational analysis. Cellular compositional and spatial metrics were computed, including the density of cells positive for each immune biomarker and a ‘proximity score’ that quantifies the relative proximity of biomarker positive cells to tumor cells. The metrics are illustrated for CD8+ cells using four NSCLC ROIs, where dots indicate the location of individual cells. The metrics across biomarkers were combined to identify distinct ‘TiME spatial factors’ through principal component analysis. Associations were then assessed between the TiME spatial factors and tumor, genomic, and clinical variables. NSCLC: non-small cell lung cancer, BRCA: breast cancer, CRC: colorectal cancer, BLAD: bladder cancer, EGC: esophagogastric cancer, MEL: melanoma, PANC: pancreatic cancer, HNC: head and neck cancer, ENDO: endometrial cancer, OVCA: ovarian cancer, RCC: renal cell carcinoma, PCa: prostate cancer, HBC: hepatobiliary cancer, SCLC: small cell lung cancer, PC: principal component.

## Results

### Pan-cancer spatial atlas of the tumor immune microenvironment

The mIF assay, named ImmunoProfile, was performed prospectively in a CLIA-certified laboratory on tumor tissue specimens from consenting patients over a three-year period at the Dana-Farber Cancer Institute^12^. The cohort includes 14 major cancer types as organized by OncoTree^13^ diagnosis codes (**Fig. 1a**). Among these, 10 cancer types have over 90 cases each. At the time of specimen collection, 850 (42%) cases were low stage (stage II or below), 1120 (55%) cases were high stage (stage III or above), and 49 (2.4%) were of unknown stage. For each case, a mIF whole slide image (WSI) was generated at 20X resolution with staining performed on a section from a formalin-fixed, paraffin-embedded (FFPE) tissue specimen. Intratumoral regions of interest (ROIs) (**Fig. 1b**) were selected by trained scientists and pathologists for each mIF WSI based on a standardized protocol. For each ROI, the x-y coordinates of cells positive for each biomarker were extracted using inForm software (Akoya Biosciences, Marlborough, MA, USA) in conjunction with custom scripts and approval by a pathologist. Further details on the assay and processing scripts are contained in the Methods. Genetic profiling was available for 1275 (63%) cases via a targeted next-generation sequencing assay performed as part of the Profile program^14,15^. In total, the analyzed cohort contains 39.4 million detected cells across 2,019 unique patients (**Supplementary Table 1**). Representing an unselected patient population with standard-of-care clinical histories, the cohort provides a unique opportunity to describe key aspects of the TiME in a real-world clinical setting.

### Identifying TiME spatial factors

We first sought to describe the major patterns of TiME variation in the cohort by generating quantitative metrics of the cellular composition and spatial organization for each case. For cell composition, we use the density of cells positive for each immune biomarker (CD8, FOXP3, PD-1, PD-L1) within the annotated intratumoral ROIs, and the PD-L1 tumor proportion score (TPS) as a tumor-specific metric. When computing cell density for the immune biomarkers, cells positive for the biomarker and negative for the tumor marker were included. To quantify spatial organization, we derived a spatial metric that measures the relative proximity of a given cell type to tumor cells in an ROI while controlling for the density of the cell type (**Fig. 1b**). This metric quantifies the likelihood that two cells are in close proximity within a ROI relative to a random distribution, calculated using a formulation based on the G-cross function^16,17^ (see Methods). We refer to this metric as the ‘proximity score’. In contrast to common spatial metrics^18^ like nearest neighbor distance, the proximity score is designed to have low correlation with cell density to enable insights into differences between spatial structure versus overall composition. We compute the proximity score separately for cells positive for each immune biomarker (CD8, FOXP3, PD-1, and PD-L1) in reference to tumor cells.

With the goal of identifying distinct patterns of variation across the pan-cancer ImmunoProfile cohort, we first assessed the pairwise correlations between all of the cell density and proximity score metrics. We observe low correlations between the proximity scores and cell densities as intended (**Supplementary Fig. 1**), but moderate correlations exist across the different biomarkers within each metric. For instance, tumors with high CD8+ cell densities also tended to have higher FOXP3+ cell densities (Spearman correlation of 0.60 (95% CI: 0.56, 0.63)). Similarly, tumors with higher CD8 proximity scores tended to have higher FOXP3 proximity scores (Spearman correlation of 0.47 (95% CI: 0.44, 0.51)). To account for these correlations, we performed principal component analysis (PCA) decomposition. PCA identifies new uncorrelated variables (principal components (PCs)) that are linear combinations of the original variables, such that the first PC captures the maximum variance across the dataset, the second captures the next highest variance orthogonal to the first, and so on.

The results of the PCA decomposition for the 2,019-patient ImmunoProfile cohort are illustrated in **Fig. 2**. The number of PCs was chosen to approximate 90% variance explained across the cohort, resulting in five PCs (89% total variance explained). **Fig. 2a** displays the variance explained and weights for these PCs, where weights above one standard deviation (s.d.) are highlighted in green and weights below negative one s.d. are highlighted in red. The first PC contains high weights for each of the immune biomarker cell densities and TPS. Tumors that score high for this “Immune Enrichment” factor demonstrate an inflamed (“hot”) phenotype, with low scoring cases indicating a “cold” phenotype. The second PC has high weights for the proximity scores across the immune biomarkers. Tumors scoring high for this “Immune-Tumor Mixing” factor demonstrate a spatially “mixed” phenotype, whereas low scoring cases indicate an “excluded” phenotype. The third PC indicates an “Immune Evasion” signature. Tumors scoring high for this factor have high TPS, high PD-L1+ cell proximity to tumor cells, and low CD8+ cell density. The fourth PC quantifies differences between the FOXP3 and CD8 proximity scores (“Treg/Cytotoxic-Tumor Mixing”) with higher values indicating relatively closer proximity of FOXP3+ cells to tumor cells than CD8+ cells to tumor cells. Analogously, PC5 largely quantifies differences between the FOXP3 and CD8 cell densities (“Treg/Cytotoxic Enrichment”). A moderately positive weight for the PD-1 proximity score is also present in PC5. **Fig. 2b** contains example ROIs scoring low and high for each of the five PCs, which we term ‘TiME spatial factors’.

**Figure 2:**
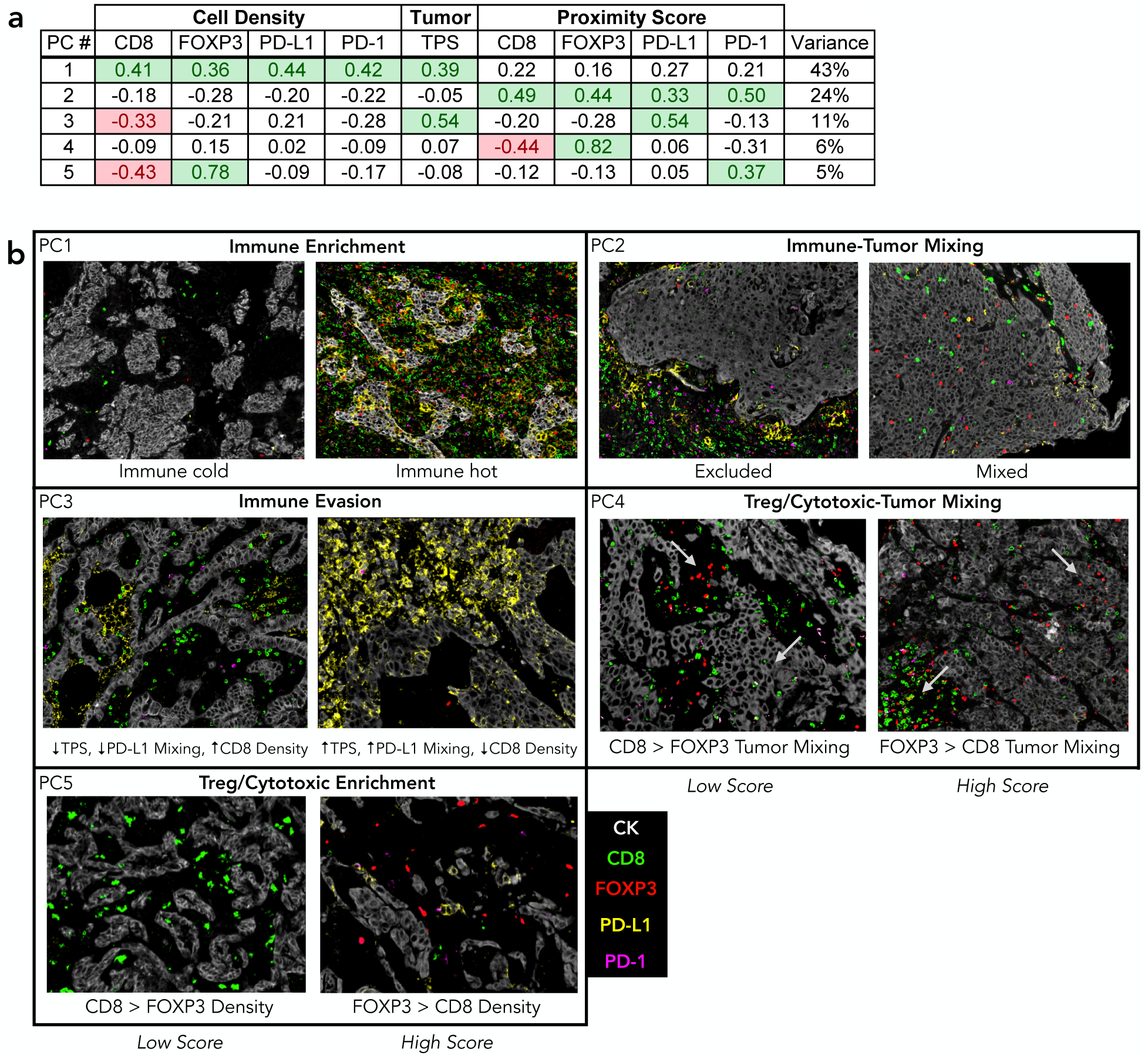
Spatial factors of the tumor immune microenvironment. a) PCA weights and corresponding attributable variance across the cohort. b) Illustration of the TiME spatial factors. mIF image patches from NSCLC ROIs scoring high and low for each factor are displayed. PC: principal component, TPS: PD-L1 tumor proportion score.

### Association of TiME spatial factors with cancer type and stage

As aspects of the TiME are generally thought to vary by cancer type, it is possible that different TiME spatial factors would be identified if the analysis was performed on certain cancer types rather than on the entire pan-cancer cohort. Alternatively, the TiME spatial factors themselves may be conserved, but the distribution of the factors may differ by cancer type (e.g., a higher proportion of ‘hotter’ tumors in some cancers). These considerations also extend to cancer stage, where high-stage tumors may reflect a failure of the immune system to control tumor growth.

To assess the possibility that different TiME spatial factors would be identified for different case subsets, we performed PCA separately for cancer types with over 200 cases (NSCLC, breast cancer, CRC), cases from the remaining cancer types, low stage cases, high stage cases, and cases with genomic profiling available. We observe similar PCA weights and variance percentages for each of these subsets (**Supplementary Fig. 2**). The correlation (*r*) in PCA weights between each subset and the entire cohort ranged from 0.89-0.99 (**Supplementary Fig. 3**). These results suggest that the five TiME spatial factors described above are generally representative of the patterns of variation across the pan-cancer cohort.

To assess whether the distribution of the TiME spatial factors varies across case subsets, we first computed scores for the TiME spatial factors for each case by applying the PCA weights. We then fit a linear mixed model^19^ to estimate the percentage of the variance in these scores that is attributable to cancer type and stage (**Fig. 3b**). The attributable variance ranged from 4.2% (Treg/Cytotoxic-Tumor Mixing (PC4)) to 18.8% (Immune Enrichment (PC1)), indicating that that the majority (81.2-95.8%) of the variability in the TiME spatial factor scores across the cohort is not explained by cancer type or stage. The variance specifically attributable to cancer type ranged from 3.4% (Immune Evasion (PC3)) to 17.8% (Treg/Cytotoxic Enrichment (PC5)), and from 0.2% (Treg/Cytotoxic-Tumor Mixing (PC4)) to 5.5% (Immune Enrichment (PC1)) for stage.

**Figure 3:**
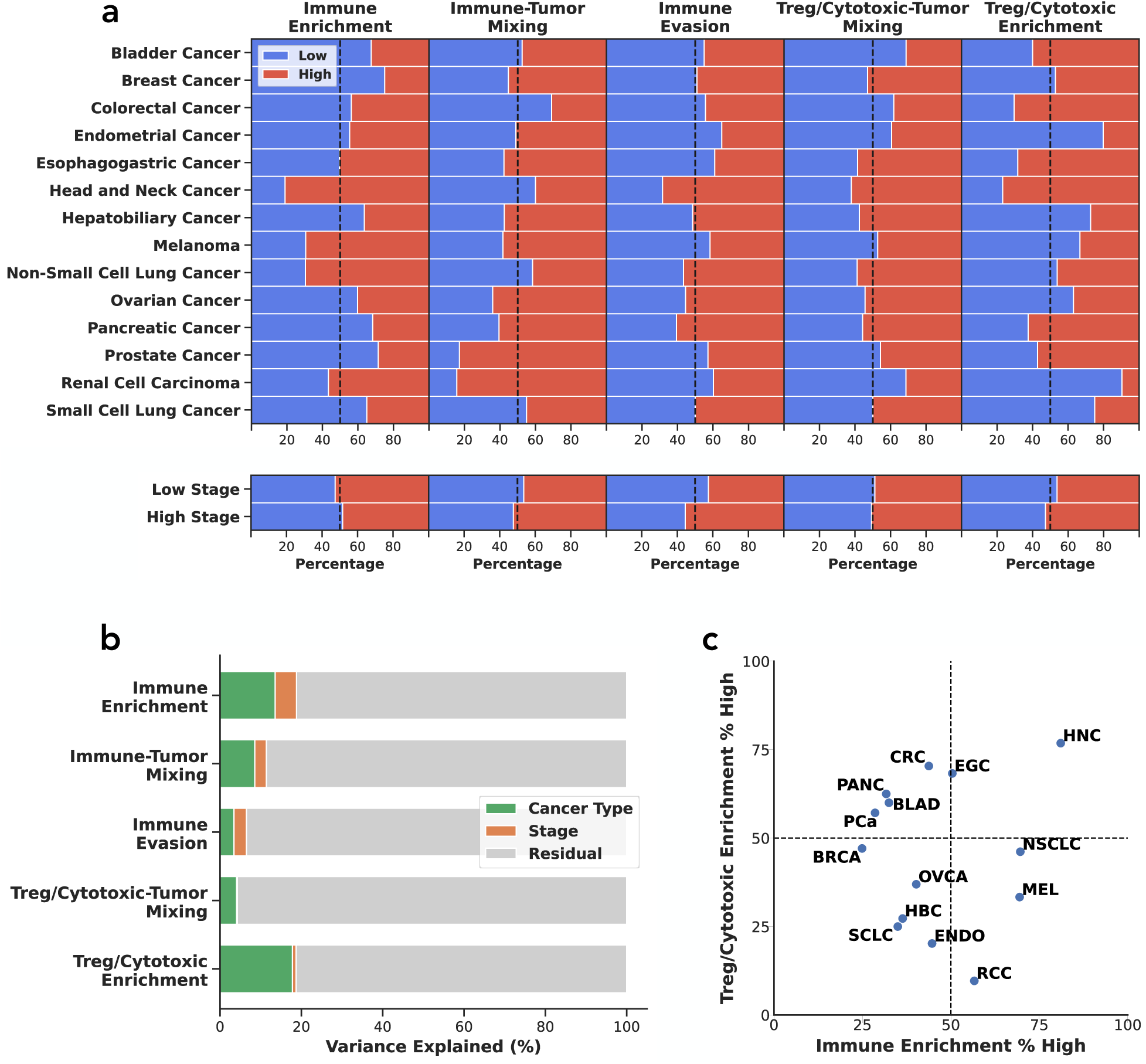
Distribution of TiME spatial factors by cancer type and stage. a) Percentage of high and low scoring cases for each factor by cancer type and stage based on a median split across the cohort. b) Variance attributable to cancer type and stage for each factor. c) Distribution of cancer types according to the Immune Enrichment and Treg/Cytotoxic Enrichment factors, which varied the most by cancer type. Axes represent the percentage of cases scoring high for the respective factor. NSCLC: non-small cell lung cancer, BRCA: breast cancer, CRC: colorectal cancer, BLAD: bladder cancer, EGC: esophagogastric cancer, MEL: melanoma, PANC: pancreatic cancer, HNC: head and neck cancer, ENDO: endometrial cancer, OVCA: ovarian cancer, RCC: renal cell carcinoma, PCa: prostate cancer, HBC: hepatobiliary cancer, SCLC: small cell lung cancer.

To visualize trends in the TiME spatial factor scores for specific cancer types/stages, we binarized the scores into high vs. low based on a median split across the cohort, followed by computing the percentage of high scores for each subset (**Fig. 3a**). For the Immune Enrichment (PC1) factor, the percentage of high scoring cases per cancer type ranged from 24.9% (breast cancer) to 81.9% (head and neck cancer). Other ‘hotter’ cancer types included non-small cell lung cancer (NSCLC), melanoma, and renal cell carcinoma (RCC); other ‘colder’ cancer types included prostate, pancreatic, and bladder. For Treg/Cytotoxic Enrichment (PC5), which varied the most by cancer type, head and neck cancer had the highest proportion of high values (i.e., more FOXP3+ cell enriched), followed by CRC and esophagogastric cancer; the lowest proportions (i.e., more CD8+ cell enriched) were observed for RCC, endometrial cancer, and small cell lung cancer (SCLC). **Fig. 3c** visualizes the Immune Enrichment and Treg/Cytotoxic Enrichment factors in combination per cancer type, where each combination of ‘hotter’/’colder’ and ‘more FOXP3 enriched/more CD8 enriched’ is represented. For cancer stage, high stage tumors had a slight trend towards higher scores for Immune Evasion (PC3) and lower scores for Immune Enrichment (PC1) (**Fig. 3a**). Altogether, while a fraction (<20%) of the variance in the TiME spatial factors is attributable to cancer type and stage, high and low values for each factor are present for each subset, further supporting the generality of the factors.

### Association of TiME spatial factors with genomic alterations

Along with cancer type and stage, the genomic phenotype of a tumor may correlate with different aspects of the TiME. We assessed associations between the TiME spatial factors and genomic alterations at three levels: somatic mutations, copy number alterations (CNAs), and tumor mutational burden (TMB). This analysis was performed for the 1275 (63%) tumors in the cohort with available genomic data. Associations were adjusted for cancer type and multiple hypothesis testing (Benjamini-Hochberg correction^20^ with a false discovery rate of 10%). For somatic mutations and CNAs, only alterations that were present in at least five cancer types and that were annotated as “oncogenic” or “likely oncogenic” by OncoKB^21,22^ were included.

**Fig. 4** summarizes the associations between the TiME spatial factors and somatic mutations. Immune Enrichment (PC1) shows the largest number of associations, including positive associations with mutations in DNA damage repair genes such as *ATR*, *BRCA1*, *MLH3*, *MSH6*, *PMS2*, and *RAD50*. The strongest negative association with Immune Enrichment was observed for *EGFR* (T=-3.61, p=0.008). The remaining factors each show an association with at least one gene: Immune-Tumor Mixing (PC2) is negatively associated with *APC* (T=-3.95, p=0.008), Immune Evasion (PC3) is positively associated with *TP53* (T=4.32, p=0.002) and negatively with *CDH1* (T=-3.22, p=0.066), Treg/Cytotoxic-Tumor Mixing (PC4) is negatively associated with *ATRX* (T=-3.13, p=0.095), and Treg/Cytotoxic Enrichment (PC5) is positively associated with *TP53* (T=3.37, p=0.077).

**Figure 4:**
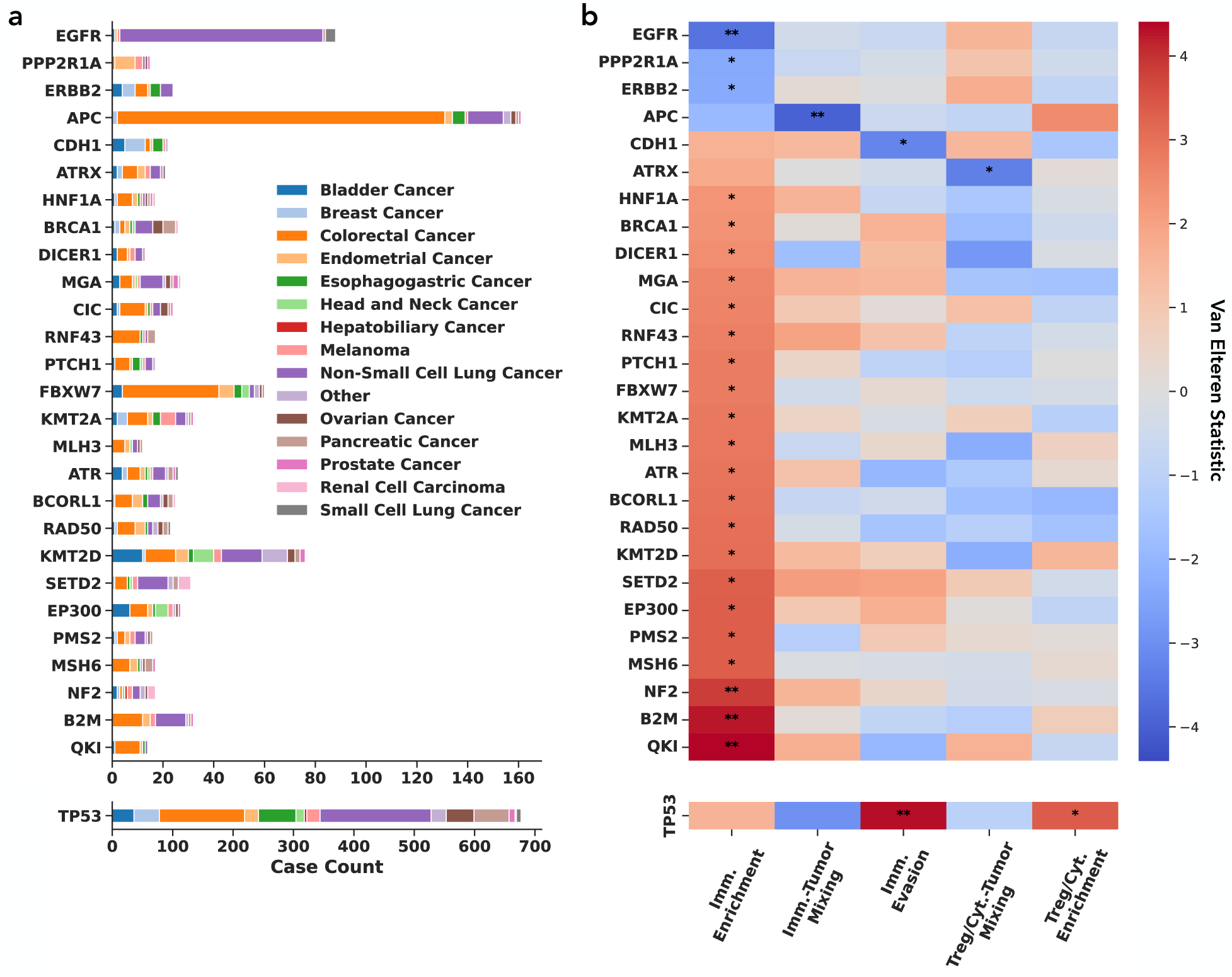
Associations between TiME spatial factors and somatic mutations. a) Counts of mutated genes by cancer type. *TP53* is shown on a separate axis given high prevalence. b) Van Elteren T-statistic per gene and TiME spatial factor. Genes with a significant association with at least one factor are displayed based on a Benjamini-Hochberg correction with a false discovery rate of 10%. (*p<0.1, **p<0.01)

Associations between the TiME spatial factors are also observed at the CNA level (**Fig. 5**). Both Immune Enrichment (PC1) and Immune Evasion (PC3) are associated with multiple CNA events. Gains in arm 9p24.1 genes *CD274*, *JAK2*, and *PDCD1LG2* had some of the strongest positive associations with Immune Enrichment (T=2.54, 2.30, 2.68, p=0.044, 0.070, 0.032, respectively), while gains in 20q13.2 genes *ZNF217* and *AURKA* had the strongest negative associations (T=-6.21, -6.00, p=3.8e-8, 7.3e-8) (**Fig. 5b**). Only negative associations were observed between CNA loss events and Immune Enrichment, with loss of arm 9p21.3 genes *MTAP*, *CDKN2A* and *CDKN2B* (T=-4.85, -4.74, -4.21, p=1.3e-4, 1.8e-4, 0.001) having the strongest negative associations (**Fig. 5d**). Strong Immune Evasion associations include gains in arm 3q genes *BCL6* and *PIK3CA* (T=4.68, 4.39, p= 2.1e-4, 4.1e-4) and arm 9p24.1 genes *CD274*, *JAK2*, and *PDCD1LG2* (T=3.70, 2.99, 3.70, p=0.004, 0.040, 0.004). Immune-Tumor Mixing (PC2) is negatively associated with gains in 20q arm genes *ZNF217* and *AURKA* and 13q arm gene *CDK8* (T=-3.14, -2.92, -3.00, p=0.08, 0.08, 0.08) (**Fig. 5d**). **Supplementary Data 1** contains results for all assessed mutations and CNAs.

**Figure 5:**
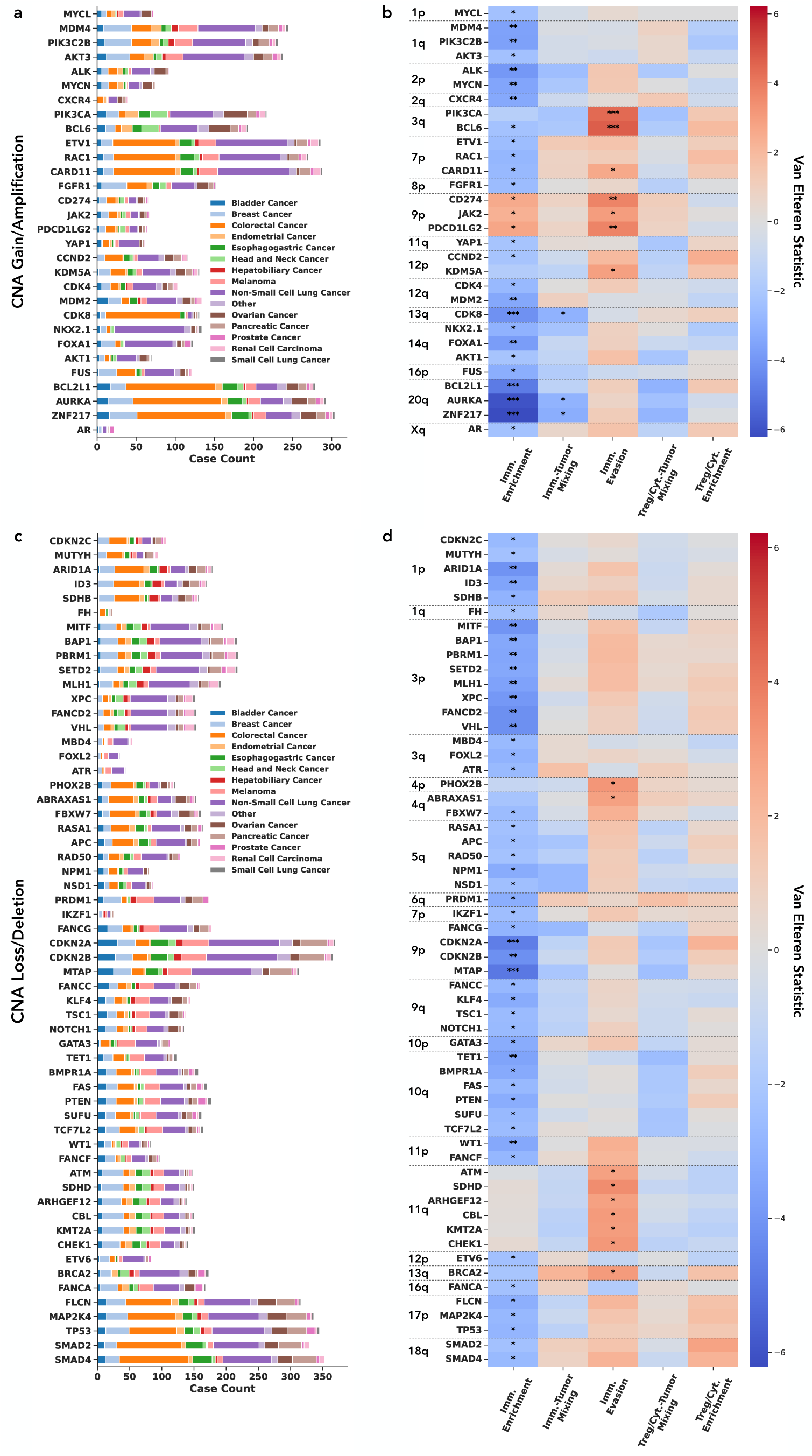
Associations between TiME spatial factors and copy number alterations. a) Counts of genes with a CNA gain. b) Van Elteren T-statistic per gene and TiME spatial factor. Genes with a significant association with at least one factor are displayed based on a Benjamini-Hochberg correction with a false discovery rate of 10%. Genes are ordered by chromosome arm. c) Counts of genes with a CNA loss. d) Van Elteren T-statistic per gene and TiME spatial factor. (*p<0.1, **p<0.01, ***p<0.001)

Finally, for TMB, we find that the Immune Enrichment (PC1) factor has a positive association with high TMB status (≥10 mutations/Mb; T=4.75, p=1.0e-5) and the Treg/Cytotoxic-Tumor Mixing (PC4) factor has a negative association (T=-2.79, p=0.013). The remaining factors do not show significant associations with TMB (Immune-Tumor Mixing (PC2): T=-2.53, p=0.80; Immune Evasion (PC3): T=1.33, p=0.31; Treg/Cytotoxic Enrichment (PC5): T=-0.92, p=0.45).

### Association of TiME spatial factors with patient prognosis

We next explored associations between the TiME spatial factors and patient overall survival (OS). **Fig. 6a** displays the concordance index (C-index) for each factor across the entire cohort (adjusted for cancer type and stage), per cancer type (adjusted for stage), and per stage (adjusted for cancer type). C-index values above 0.50 indicate positive correlation with survival, values below 0.50 indicate negative correlation with survival, and a value of 0.50 indicates a chance result. The most prognostic factor across the cohort is Immune Evasion (PC3) with a C-index of 0.40 (p=3.1e-7), signifying that higher scores for this factor are associated with reduced survival. This trend is consistent across cancer types and stages. For a sense of magnitude, tumors that score high for Immune Evasion have a 67% increased risk compared to those that score low (hazard ratio (HR) of 1.67 (95% CI: 1.41, 1.99) when binarizing into ‘high’ vs. ‘low’ as in **Fig. 3a**). The second most prognostic factor is Immune Enrichment (PC1) with a C-index of 0.57 (p=2.4e-6; HR of 0.70 (95% CI: 0.59, 0.83) when binarized into ‘high’ vs. ‘low’), indicating a positive association with survival. This association appears to be largely driven by high stage tumors, as an association is not observed for low stage tumors (**Fig. 6a**). The remaining factors (Immune-Tumor Mixing (PC2), Treg/Cytotoxic-Tumor Mixing (PC4), Treg/Cytotoxic Enrichment (PC5)) all have negative associations with survival at the cohort level. For individual cancer types, there is some variation in the direction and magnitude of the C-index per factor, but all significant associations have consistent directions across the cohort.

**Figure 6:**
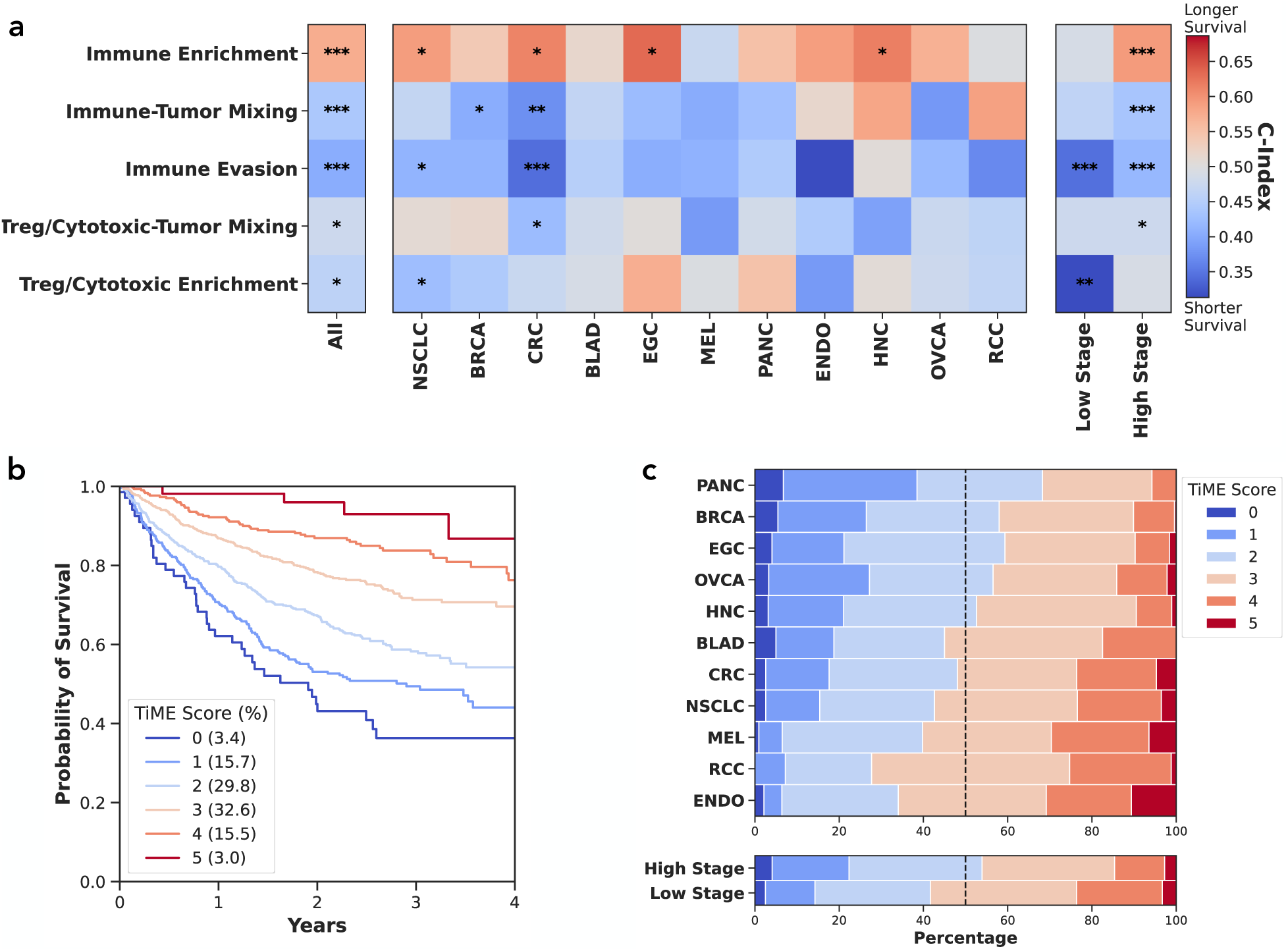
Associations between TiME spatial factors and prognosis. a) Concordance index (C-index) between factors and patient overall survival for the entire cohort, per cancer type, and per stage. Values above 0.50 indicate positive correlation with survival, values below 0.50 indicate negative correlation with survival, and a value of 0.50 indicates a chance result. Cancer types with at least 80 cases are displayed. b) Kaplan-Meier curve based on the TiME Score across the cohort. The TiME Score represents the number of factors that are in the positive direction for survival for a given case. c) Distribution of TiME Score by cancer type and stage. Low and high scores are present for each subset. NSCLC: non-small cell lung cancer, BRCA: breast cancer, CRC: colorectal cancer, BLAD: bladder cancer, EGC: esophagogastric cancer, MEL: melanoma, PANC: pancreatic cancer, HNC: head and neck cancer, ENDO: endometrial cancer, OVCA: ovarian cancer, RCC: renal cell carcinoma. (*p<0.1, **p<0.01, ***p<0.001)

While our goal was to elucidate TiME variability rather than maximize prognostic accuracy per se, it is useful to consider how the combination of the TiME spatial factors relates to prognostic stratification. To do so in an intuitive manner, we constructed a composite “TiME score” for each tumor based on whether it scores high or low for each factor. The composite score is defined as the number of factors that align positively with survival time based on the associations calculated across the cohort. As shown in **Fig. 6b**, we find that this TiME score, ranging from 0 to 5, monotonically stratifies patients by risk (p=4.3e-17, p-value adjusted for cancer type and stage). Importantly, this stratification does not simply result from differences in TiME scores across cancer types or stages, as both high and low scores are observed for each subgroup (**Fig. 6c**) and the stratification is evident even within individual cancer types (**Supplementary Fig. 4).** Among cancer types, head and neck cancers had the highest proportion of favorable TiME scores and endometrial cancer had the lowest.

## Discussion

We characterized tumor-immune topographic organization across a real-world, pan-cancer cohort of 2,019 patients. By combining density and spatial metrics of key immune biomarkers, we identified five TiME spatial factors that are conserved across cancer types and stages. The factors provide different axes to describe tumors in a data-driven fashion. We identified trends in the distribution of the factors across clinicopathological variables, including identifying associations with molecular signatures and patient prognosis.

While some TiME spatial factors and their associations align with biological intuition and prior findings, unexpected trends also emerge. Descriptors of “hot/cold” and “excluded/mixed” are commonly used in immuno-oncology^23^, and these axes emerge as the first two factors in our unsupervised decomposition analysis. This analysis results in quantitative definitions of these commonly used terms and highlights other considerations, such as including high PD-L1 TPS as a component of the immune ‘hot’ phenotype. Other intuitive factors such as the difference between cytotoxic CD8+ T-cell and regulatory FOXP3+ T cell enrichment also emerge from our analysis. The ‘Immune Evasion’ (PC3) factor provides a nuanced means to describe the TiME, wherein tumors scoring high for this factor have high PD-L1 TPS, high PD-L1+ cell proximity to tumor cells, and low CD8+ cell density. Studies have shown that PD-L1+ immune cells are commonly macrophages^24^ and certain tumor-associated macrophages can promote immune evasion^10,25,26^. Since the TiME spatial factors correspond to orthogonal sources of variation across the cohort, they can be used as complementary ways to describe tumors. For instance, a tumor could be ‘hot’ but also score high for Immune Evasion based on the relative differences between certain immune biomarker densities and spatial arrangements.

Unsupervised decomposition performed on different case subsets showed results largely consistent with those obtained across the entire cohort, suggesting that the identified TiME spatial factors may represent fundamental properties of tumor-immune cell interactions that are conserved across solid tumors. Furthermore, cancer type and stage accounted for <20% of the variance in the distribution of factor scores across the cohort. Nonetheless, several trends were observed. The TiME spatial factors that varied most by cancer type were Immune Enrichment (PC1) and Treg/Cytotoxic Enrichment (PC5). Head and neck cancer, NSCLC, and melanoma scored the highest for Immune Enrichment, aligning with the general understanding that these are ‘hotter’ tumors^23,27^. In contrast, the ‘coldest’ tumors were breast, prostate, and pancreatic, also aligning with prior knowledge^23,27^. In terms of the relative difference between Treg and Cytotoxic T-cell enrichment, head and neck cancer was the most Treg enriched, aligning with previous reports of a high FOXP3/CD8 ratio in this cancer type^28,29^, and RCC was the most Cytotoxic T-cell enriched.

In terms of genomic alterations, the largest number of associations was observed for the Immune Enrichment (PC1) factor. Somatic mutations in genes involved in the DNA mismatch repair (MMR) pathway – *MLH3*, *MSH6*, and *PMS2* – were associated with higher Immune Enrichment scores, consistent with the high immunogenicity of MMR-deficient tumors. Beta2-microglobulin (*B2M*) also exhibited a strong positive association with Immune Enrichment. *B2M* alterations have been shown to correlate with poor immunotherapy response^30,31^, but are also associated with MMR deficiency and high TMB^31^. *EGFR* mutant tumors demonstrated the strongest negative association with Immune Enrichment, aligning with evidence of an uninflamed phenotype^32^. Notably, lung adenocarcinomas with activating *EGFR* mutations are typically unresponsive to immunotherapy^33^. Additionally, *APC* mutations were negatively associated with the Immune-Tumor Mixing (PC2) factor, suggesting an immune-excluded phenotype. A prior study identified a negative correlation between *APC* mutation and immunotherapy response in CRC^34^, so our results may further help explain this finding. *TP53* mutations were positively associated with Immune Evasion (PC3) and Treg/Cytotoxic Enrichment (PC5), consistent with studies linking *TP53* with PD-L1 and FOXP3 expression^35–38^. Finally, *ATRX* mutations were negatively associated with the Treg/Cytotoxic-Tumor Mixing (PC4) factor, indicating a correlation with higher cytotoxic T-cell proximity to tumor cells relative to Treg proximity. This finding may align with a prior study reporting increased immunotherapy efficacy and increased T cell infiltration in *ATRX*-mutant NSCLC tumors^39^.

For copy number alterations, positive associations were found between gains in chromosome 9p24.1 genes – *CD274* (PD-L1), *JAK2*, and *PDCD1LG2* (PD-L2) – and both Immune Enrichment (PC1) and Immune Evasion (PC3). 9p24.1 amplification has been previously associated with high, but ineffective immune infiltration with high PD-L1 expression in classical Hodgkin lymphoma^40^. Our findings suggest that these associations may be more widespread. These findings also align with evidence that 9p24.1 losses are linked to immune-cold, ICI-resistant tumors, while 9p24.1 gains are associated with immune-hot, ICI-responsive tumors in head and neck cancer^41^. In addition, 9p arm-level loss, which includes *CDKN2A/B* and *MTAP*, was previously associated with impaired outcomes to immunotherapy in advanced NSCLC^42^. Of all genetic alterations and TiME spatial factors, the strongest associations were found between amplification of 20q13.2 genes *BCL2L1*, *AURKA*, and *ZNF217* and decreased Immune Enrichment (PC1). Aurora-A (AURKA) inhibitors are an emerging targeted therapy, and studies suggest that their use can enhance immune infiltration^43,44^. Finally, activating mutations of *PIK3CA* have been correlated with PD-L1 expression^45^ in CRC and improved survival^46^ for CRC patients treated with celecoxib (COX2-targeted anti-inflammatory), and gains in 3q genes have been associated with ICI response in cutaneous squamous cell carcinoma^47^. In our cohort, gains in 3q genes *PIK3CA* and *BCL6* were negatively associated with Immune Enrichment (PC1) and positively associated with Immune Evasion (PC3). Overall, we note that it can be difficult to isolate gene-level effects from what may be large region or arm-level alterations with CNA analysis, but our findings nonetheless provide insights into the association between the TiME and genetic phenotypes. Furthermore, we stratified by cancer type when assessing genomic associations and only included alterations that are present in at least five cancer types, but some associations may still be driven by skewed occurrence in specific cancers (**Fig. 4a**, **Fig. 5a,c**).

Although our primary aim was to characterize TiME variation and its biological correlates, each TiME spatial factor was associated with patient overall survival. The most prognostic factor was Immune Evasion (PC3), which showed consistently negative correlations with survival across cancer types and stages. Immune Enrichment (PC1) and Treg/Cytotoxic Enrichment (PC5) were positively and negatively correlated with survival, respectively; however, these trends varied by stage. The Immune Enrichment pattern was driven by high stage cases, whereas the Treg/Cytotoxic Enrichment pattern was driven by low stage. This may suggest that the extent of immune infiltration is prognostically important as cancer progresses, but the relative type of infiltration is most important at an early stage. For the Immune-Tumor Mixing (PC2) factor, an unexpected negative association with survival was observed. There may be several reasons why this is the case. First, this factor has positive weights for the proximity scores of all the immune biomarkers, whereas the association with survival may be biomarker specific. Indeed, relatively higher cytotoxic T-cell proximity than Treg proximity to tumor cells was positively correlated with survival, as quantified by the Treg/Cytotoxic-Tumor Mixing (PC4) factor. Second, while the Immune-Tumor Mixing (PC2) factor had the highest weights for the proximity scores, slightly negative weights were also present for the cell density metrics, which may also contribute to the negative correlation with prognosis. Altogether, the prognosis analysis highlights the potential of quantitative, image-based prognostic biomarkers, but future work is necessary to maximize this prognostic accuracy, such as using supervised machine learning techniques. For instance, our group has shown that CD8+ cell density alone is significantly prognostic in the ImmunoProfile cohort^12,48^, demonstrating similar prognostic accuracy (C-index of 0.61) as the Immune Evasion factor (C-index of 0.60) identified in this study which weighs CD8+ cell density and PD-L1 metrics in an unsupervised fashion.

In contrast to prior pan-cancer tumor subtyping analyses using cohorts such as TCGA^9,11^ and CPTAC^8^, our analysis captures single-cell, spatial aspects of the TiME through a prospectively performed mIF assay. Given differences in methodological approaches and aspects of the TiME being measured, it is challenging to directly compare our TiME spatial factors to these previously reported immune subtypes. The most relevant comparison arguably lies in the existence of hot versus cold descriptors across the different schemes, such as the Immune Enrichment (PC1) factor in our work, inflammatory and lymphocyte-depleted subtypes in Thorsson et al.^9^, immune-enriched and immune-depleted subtypes in Bagaev et al.^11^, and the CD8+/IFNG+ and CD8-/IFNG-subtypes in Petralia et al.^8^

While large-scale, pan-cancer mIF studies are currently lacking, more highly plexed assays have been performed in smaller studies that image tens of biomarkers compared to five in the ImmunoProfile cohort. Thus, our analysis is inherently confined to the studied biomarkers; however, these biomarkers (i.e., CD8, FOXP3, PD-1, PD-L1) have been implicated in many aspects of tumor immunity^49–51^. An additional challenge, but also a strength, of the studied cohort is its heterogeneity. Representing an unselected clinical population, the cohort consists of patients undergoing varying treatment regimens across different cancer types. Nonetheless, our core findings were robust across cancer types and stages, enabling the identification of overarching associations as well as cancer specific trends. The prognosis analysis should especially be interpreted in the context of the current standard of care, where treatment-specific effects are challenging to estimate in pan-cancer settings and may shift as new approaches emerge. Finally, our methodological approach was designed to disentangle immune biomarker densities from their tumor proximity and result in interpretable factors, but higher-order levels of spatial organization would also be fruitful to analyze in large cohorts, including non-intratumoral regions, intratumoral heterogeneity, and whole slide image (WSI) analysis. While the ROI sampling strategy utilized here leverages pathologist expertise and reduces computational costs, WSI-level analysis or automated ROI sampling strategies may ultimately limit potential sampling biases and enhance reproducibility across clinical sites.

Understanding the spatial organization of the TiME and its relation to genetic profiles and patient outcomes is a critical path forward in precision oncology. Our analysis and accompanying database of 39.4 million spatially-resolved cells is designed to accelerate progress in this direction.

## Methods

### Ethics statement

All patients in the ImmunoProfile cohort provided consent under an institutional research protocol (DFCI 11-104, 17-000, or 20-000).

### ImmunoProfile assay

For each case, a formalin-fixed, paraffin-embedded (FFPE) tissue block from either a core needle biopsy or resection was sectioned to generate a hematoxylin and eosin (H&E) stained slide and an unstained slide. The H&E slide was digitized and reviewed by a pathologist to confirm the diagnosis and assess tumor content. The unstained slide was used for automated multiplex immunofluorescence (mIF) staining using a BOND Rx® system (Leica Biosystems, Danvers, MA, USA). The mIF panel included five markers: CD8, FOXP3, PD-1, PD-L1, and a tumor-specific marker (cytokeratin for epithelial cancers, PAX8 for renal cell carcinomas, and SOX10 for melanomas). DAPI was used as a nuclear counterstain. The mIF-stained slides were imaged using a PhenoImager® (Akoya Biosciences, Marlborough, MA, USA) scanner at 20X magnification.

Regions of interest (ROIs) representative of the overall tumor microenvironment within each mIF whole slide image (WSI) were selected by a technician scientist and approved by a pathologist. Between three and six ROIs were selected for each slide depending on the amount of available tissue. For each ROI, the inForm software (Akoya Biosciences, Marlborough, MA, USA) was used to perform spectral unmixing, cell segmentation, and quantification of marker intensity for each cell. The scientist then selected cells that were positive and negative for each biomarker in the ROI based on their assessment of staining patterns, which were used to train a random forest algorithm in the inForm software. The trained algorithm would then output positive and negative biomarker labels for the remainder of the cells in the ROI. The scientist also annotated the tumor border if present for each ROI, specifically segmenting intratumor, tumor stromal interface, and stroma regions using an internally developed GNU Image Manipulation Program. Custom algorithms were employed to calculate the PD-L1 Tumor Proportion Score (TPS) and the densities of non-tumor cells positive for each immune biomarker. These calculations were performed across the union of intra-tumor annotated regions for a given case. The annotation tools and algorithms can be found at: https://github.com/dfci/pythologist. All of these steps and the subsequent results were reviewed and approved by a pathologist using a digital sign-out tool.

The ImmunoProfile assay and its workflow were validated within the CLIA-certified molecular diagnostics laboratory at the Brigham & Women’s Hospital in Boston (CLIA license #22D2156930). Quality control testing resulted in the assay’s full clinical validation in compliance with CLIA standards. This validation included demonstrating high concordance between the assay and immunohistochemistry (IHC) and consistent inter-run reproducibility over time. Further details of the workflow and validation can be found in ref. 48.

### Clinical cohort

Patients in the ImmunoProfile cohort were enrolled at the Dana-Farber Cancer Institute (DFCI) in Boston, MA between November 2018 to January 2022. Upon patient consent to one of the institutional research protocols listed above, the decision to order the ImmunoProfile assay was made by the patient’s clinical team. All solid cancers besides sarcomas, pediatric tumors, and neuro-oncology cases were eligible for the assay. Data for each patient, including cancer type, vital status/date of death, systemic therapies received at DFCI, specimen date, and cancer stage, were curated from medical records and internal databases with a retrieval date of August 2022. To ensure accuracy, clinical domain experts manually reviewed each case against the patient’s medical record, including confirming that the reported stage reflected the disease status at the time of specimen collection. From the original cohort of 2,023 cases, four cases were excluded from the analysis because of insufficient numbers of cells to reliably compute spatial metrics, resulting in the final number of 2,019 cases (each from a unique patient). The analyzed cohort is summarized in **Supplementary Table 1**. As a real-world clinical cohort, patients underwent a variety of treatment regimens, including 466 (23%) patients having received immunotherapy. Of these 466 patients, 234 started immunotherapy after the date of the tissue specimen used for ImmunoProfile, 131 started immunotherapy before, and 101 had an unknown immunotherapy start date. A total of 631 (31%) patients experienced a death event, with a median of 326 days (IQR: [137, 587]) from tissue sample collection to death. For censored patients, the median follow-up time was 923 days (IQR: [684, 1164]). The mean age was 63.7 years (IQR: [56.2, 72.6]). The distribution of patient sex was 1213 (56%) female patients and 946 (44%) male patients. The distribution of self-reported race was as follows: 1838 (91.0%) white, 59 (2.9%) Asian, 58 (2.9%) Black, 30 (1.5%) Other, 4 (0.2%) Mixed, 1 (<0.1%) Native Hawaiian or Other Islander, 29 (1.4%) Unknown/Declined.

### Proximity score metric

The proximity score metric was formulated based on the G-cross function^9,16,17^. The G-cross function between cell types *A* and *B*, denoted as *G*_*A,B*_(*r*), represents the cumulative probability that a cell of type *B* is within a radius *r* from a cell of type *A*. We computed this function separately for each immune biomarker with respect to tumor cells for a given ROI. Explicitly, cell type *A* corresponded to cells positive for the tumor marker and cell type *B* corresponded to cells negative for the tumor marker and positive for a specific immune biomarker (CD8, FOXP3, PD-1, or PD-L1). The G-cross curve was then summarized into a single value using the area under the curve (AUC) up to a radius of 100 pixels (approximately 50 microns). The proximity score was then formulated as the ratio of this empirical AUC to a theoretical AUC that assumes random distribution of cells. Mathematically, the theoretical G-cross curve takes the form of *G_A,B_*(*r*) = 1 – *e*^λ*B*π^*^r^*^2^ where λ*_B_* is the density of cell type B in the ROI. Higher proximity scores thus indicate relatively closer immune cell-tumor cell proximity than would be expected based on the density of the immune cells alone. The proximity scores were calculated for ROIs for which at least 90% of cells fell in the segmented intratumor region and there were at least 20 cells of each type involved in the calculation. These criteria were employed to help mitigate potential confounders introduced by non-tumor regions and to standardize the calculations across cases, and because the relative proximity of cells is ill-defined/noisy with low cell counts. A case-level value was computed as the mean across ROIs. Both the density and proximity score metrics were performed at an individual biomarker level rather than combinations of biomarkers (e.g., CD8+PD1+) because of the relatively low frequency of non-tumor cells positive for more than one biomarker, which would especially complicate the spatial analysis and hinder interpretability of the PCA. The R package *spatstat*^52^ was used to compute the G-cross function.

### Principal component analysis

The input variables to the PCA consisted of the cell density and proximity score for each immune biomarker (CD8, FOXP3, PD-1, PD-L1) and PD-L1 TPS. Each variable was normalized using a percentile ranking across the cohort to account for different scales across the variables and to emphasize the relative ranking across cases rather than the absolute magnitude. Missing proximity score values due to the cell count threshold described above were imputed using the median across the cohort. The rationale for this imputation is that the relative spatial arrangement of the immune cells within the ROI are neither closer nor farther away to the tumor than average, but rather undefined due to low counts, which is already accounted for in the cell density metric. The percentage of cases with imputed proximity scores was 15%, 30%, 28%, and 26% for CD8, FOXP3, PD-1, and PD-L1, respectively.

The PCA was performed across the entire cohort with subgroup analysis to assess consistency across specific case subsets (**Supplementary Fig. 2**). The number of principal components retained was chosen based on the value that was closest to explaining 90% of the variance, which resulted in five components for the cohort (89% variance explained). To compare the PCs computed over different case subsets, a matching algorithm was first performed to map PC numbers across the subsets. Given two sets of PC weights, the absolute value in the correlation (*r*) between each pair of PCs between set 1 and set 2 was first computed. Starting with the first PC in set 1, the corresponding matching number in set 2 was assigned based on the highest correlation. This number was then removed from the list of possible matches and the process proceeded until a match was assigned for each PC in set 1. **Supplementary Fig. 2** displays the resulting ordering and PC weights for different case subsets in comparison to the PCA fit across the cohort. **Supplementary Fig. 3** displays the aggregate correlation in the PC weights between all pairs of subsets based. The correlation (*r*) is computed based on concatenating the weights of the first five PCs for each subset after the described alignment process.

### Associations with cancer type and stage

The fraction of the variance in the TiME factor scores that is attributable to cancer type and stage was estimated using the variancePartition software^19^, which fits a linear mixed model. Cancer types with less than 20 cases were grouped as ‘Other’ for this analysis and stage was binarized into low (0, I, II) or high (III, IV). Variations in the TiME factors for specific cancer types and stages were then visualized by binarizing the scores for each factor based on a median split across the cohort.

### Associations with genomic alterations

1275 ImmunoProfile cases were sequenced using OncoPanel (OP)^14,15^, a custom hybrid-capture sequencing assay which targets exons of 304–447 genes across three panel versions (v1 N=22 cases, v2 N=97 cases, v3 N=1156 cases). TMB was calculated for the 1156 OPv3 cases as the number of coding somatic mutations that occur per megabase of exonic sequence data across all genes on the panel. Cancer type stratified van Elteren^53^ tests were run to compare TiME spatial factor score distributions with genomic alterations using the *npsm*^54^ package in R. For TMB, strata included cancer types which represented at least 10% of all cases with high TMB (≥10 mutations/Mb), which included NSCLC, CRC, and melanoma with the remaining cancer types grouped as ‘Other’. Associations were assessed using the binarized high TMB status. For mutation and CNA gain/loss analyses, strata included cancer types which represented at least 5% of all alteration events for the alteration category, which consisted of the following; mutation events: bladder, CRC, endometrial, esophagogastric, melanoma, NSCLC, and pancreatic; CNA gains: breast, CRC, esophagogastric, melanoma, NSCLC, ovarian; CNA losses: breast, CRC, esophagogastric, melanoma, NSCLC, ovarian, pancreatic; with remaining cancer types grouped as ‘Other’ in each scenario. The list of mutations/CNA events included in the analysis was curated by starting with the genes present in OPv3, followed by annotation using OncoKB-Annotator (https://github.com/oncokb/oncokb-annotator) and filtering for alterations annotated as “oncogenic” or “likely oncogenic” ^21,22^. To focus on pan-cancer effects, genes were further filtered to include only those with an alteration event in at least 5 different cancer types. Rare events (less than ten total alterations among all cases) were also excluded. For a given gene, all cases for which the gene is part of the OncoPanel version used for the case were included in the analysis (i.e., if a gene was in versions 1-3, all cases were included, whereas if it was only present in v3, only v3 cases were included). The full list of all tested genes, their panel version, and results are contained in **Supplementary Data 1.**

### Associations with patient prognosis

Associations between the TiME spatial factors and patient prognosis were assessed using Cox proportional hazard (CoxPH) models^55^ with cancer type and stage (low vs. high) as stratification variables. Models were fit separately for each TiME spatial factor using the entire cohort, for each cancer type with at least 80 cases, and for high and low stage. To explore multivariate prognostication, a composite TiME score was calculated for each case that equaled the number of TiME factors with binarized values corresponding to better prognosis. Binarized values were calculated using a median split across the cohort and the direction of better prognosis was based on the univariate results. For instance, if a case scores ‘high’ for Immune Enrichment, the case will be given a score of 1 for this factor since it was positively associated with survival. Conversely, a high value for Immune Evasion would correspond to a score of 0 as this factor is negatively associated with survival. The composite TiME score is the sum across the five factors, yielding a score between 0 and 5.

### Additional statistical considerations

For the genomics and prognosis analyses, the Benjamini-Hochberg method^20^ was performed to correct for multiple comparisons with a false discovery rate of 10% for significance. All p-values are two sided unless otherwise noted. Statistical methods pertaining to specific analyses are described in their respective sections above.

## Resource availability

### Lead contact

Further information and requests for resources and reagents should be directed to and will be fulfilled by the lead contact, William Lotter.

### Materials availability

This study did not generate new unique reagents.

### Data and code availability

- The single-cell database with biomarker expression, genomic, and clinical annotations is deposited at [url upon publication] and are publicly available as of the date of publication.
- All original code has been deposited on Github and is publicly available at [url upon publication] as of the date of publication.

## Figures and Tables

**Supplementary Table 1:**
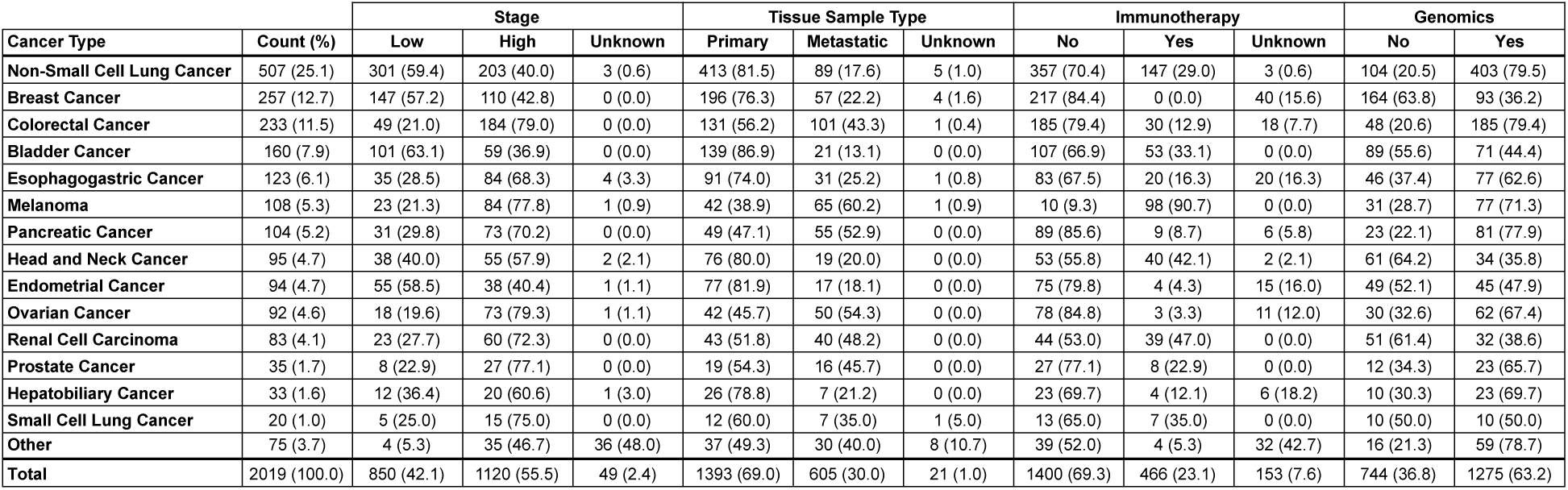
Distribution of cases in cohort. Tissue sample type indicates whether the specimen represents a primary or metastatic site. Genomics indicates the availability of genomic profiling. Immunotherapy indicates whether the patient was treated with immunotherapy either before or after the specimen date. Cancer types with less than 20 cases were grouped as ‘Other’ and include the following: Cancer of Unknown Primary (17 cases), Cervical Cancer (12), Thyroid Cancer (11), Anal Cancer (8), Gastrointestinal Neuroendocrine Tumor (5), Small Bowel Cancer (5), Salivary Gland Cancer (5), Non-Melanoma Skin Cancer (4), Mesothelioma (3), Ampullary Cancer (3), Vaginal Cancer (1), Thymic Carcinoma (1).

**Supplementary Figure 1:**
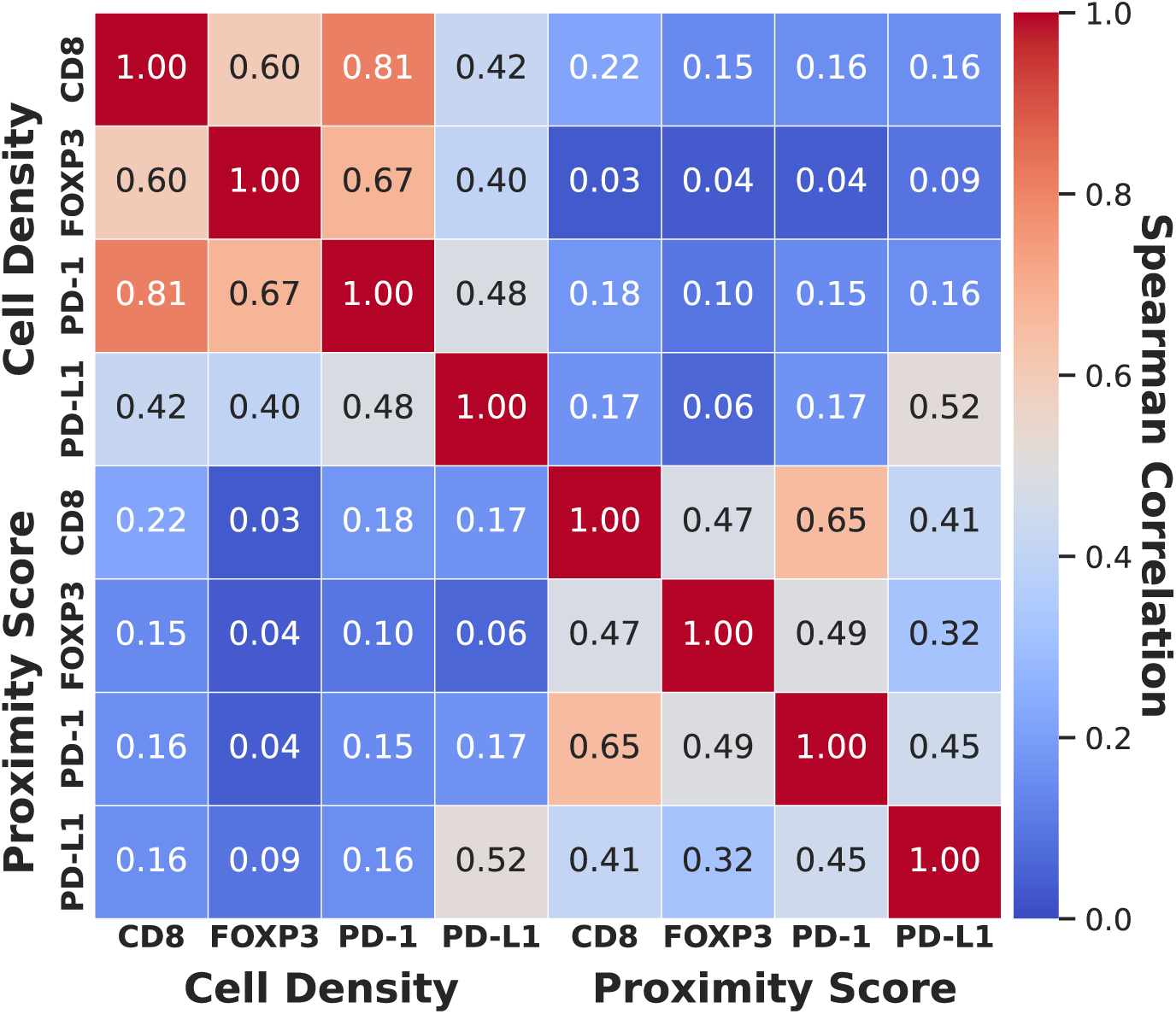
Correlations between the cell density and proximity score metrics across biomarkers. Values represent the Spearman correlation computed across the cohort. Low correlations are generally observed between the cell density and proximity score metrics for the same biomarker, and moderate to high correlations are observed between different biomarkers for the same metric.

**Supplementary Figure 2:**
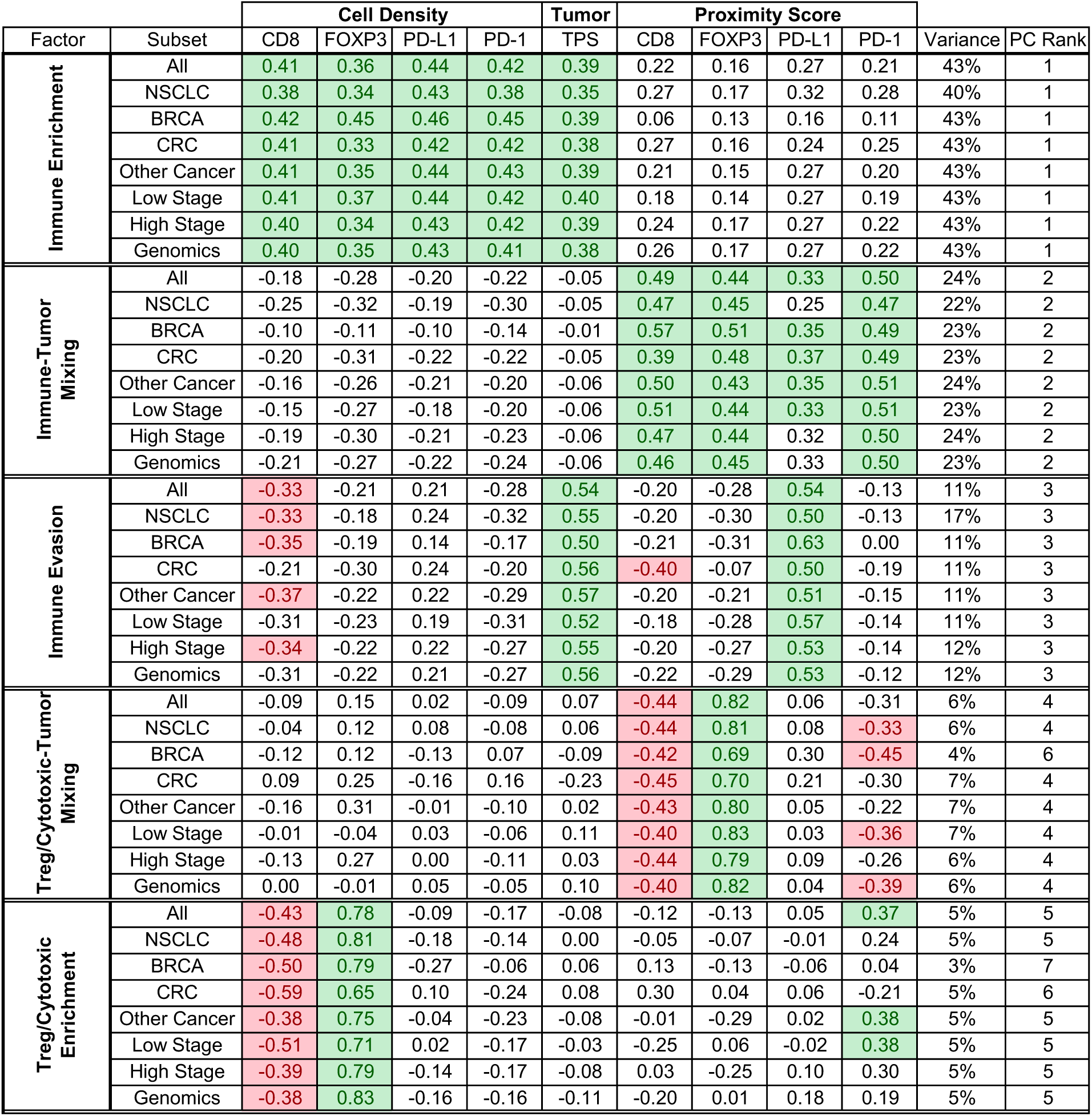
Results of PCA performed on different case subsets. PC rank indicates the original order of the PC in terms of explained variance for the subset. High consistency is observed across the case subsets in terms of both the PC weights and variance explained. ‘Other Cancer’ includes cases from any cancer other than NSCLC, BRCA, and CRC. Genomics includes cases for which genomic profiling is available. NSCLC: non-small cell lung cancer, BRCA: breast cancer, CRC: colorectal cancer, PC: principal component, PCA: principal component analysis.

**Supplementary Figure 3:**
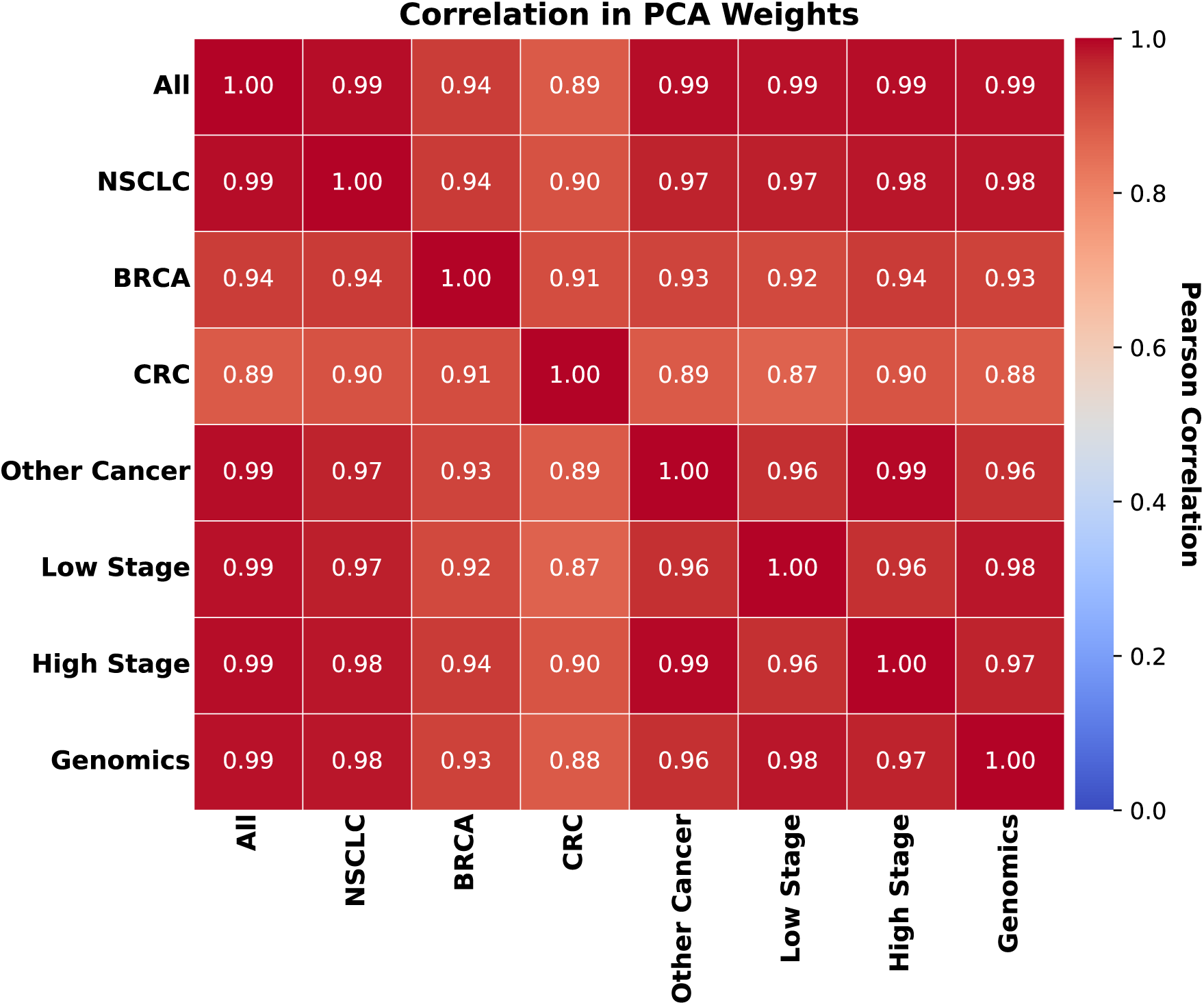
Correlation in principal component weights between different case subsets. The correlation is computed between the five PCs representing the TiME spatial factors described in the text. High correlation is observed between each pair. ‘Other Cancer’ includes cases from any cancer other than NSCLC, BRCA, and CRC. Genomics includes cases for which genomic profiling is available. NSCLC: non-small cell lung cancer, BRCA: breast cancer, CRC: colorectal cancer, PC: principal component, PCA: principal component analysis.

**Supplementary Figure 4:**
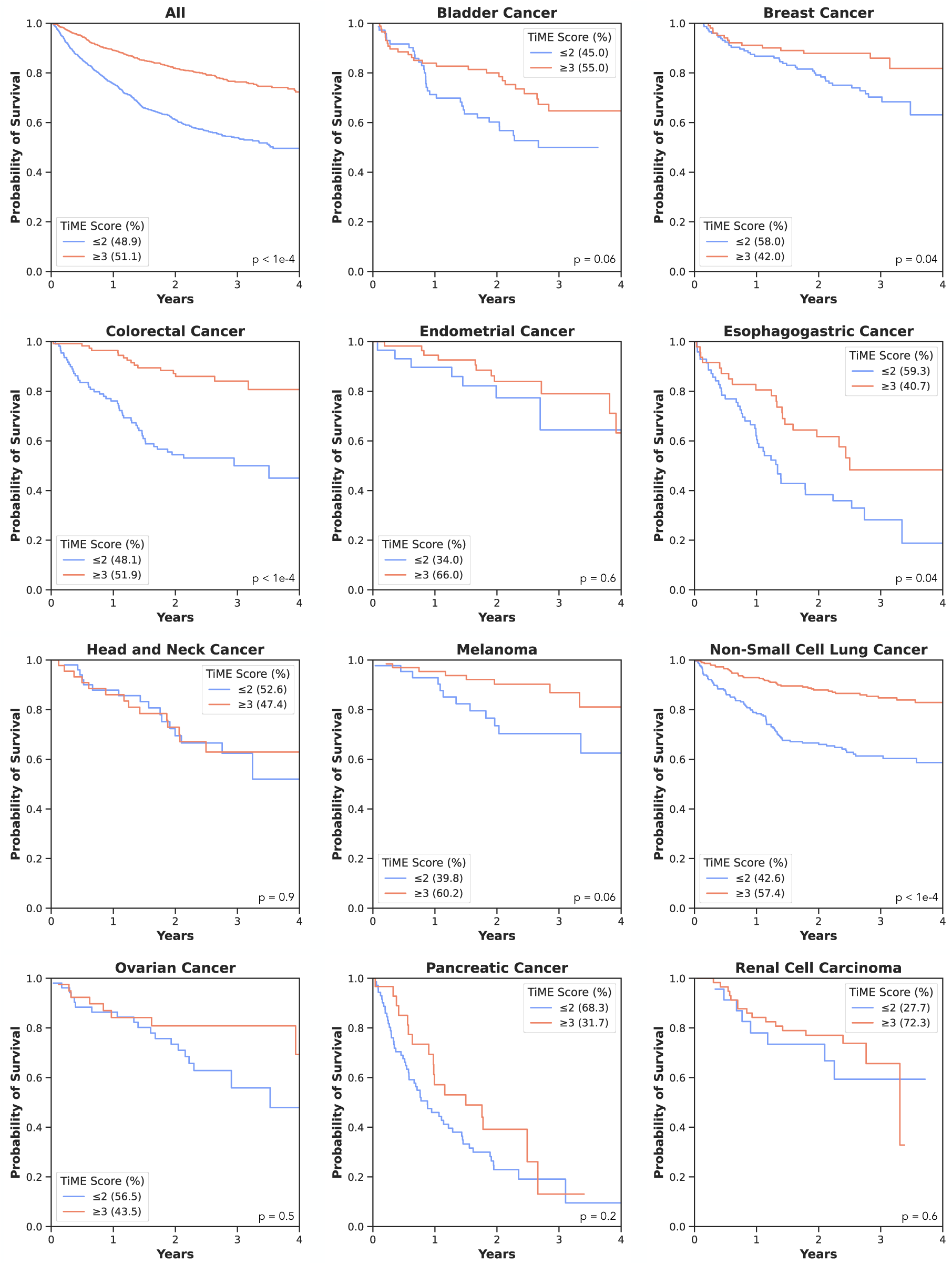
Kaplan-Meier curves by TiME Score per cancer type. The TiME Score is formulated based on the entire cohort and is displayed per cancer type. Scores are split into low (≤2) or high (≥3) due to sample sizes. Two-sided p-values are computed via the log-rank test and are Benjamini-Hochberg corrected for multiple comparisons.

